# Distinct shifts in site-specific glycosylation pattern of SARS-CoV-2 spike proteins associated with arising mutations in the D614G and Alpha variants

**DOI:** 10.1101/2021.07.21.453140

**Authors:** Chu-Wei Kuo, Tzu-Jing Yang, Yu-Chun Chien, Pei-Yu Yu, Shang-Te Danny Hsu, Kay-Hooi Khoo

## Abstract

Extensive glycosylation of the spike protein of SARS-CoV-2 virus not only shields the major part of it from host immune responses, but glycans at specific sites also act on its conformation dynamics and contribute to efficient host receptor binding, and hence infectivity. As variants of concern arise during the course of the COVID-19 pandemic, it is unclear if mutations accumulated within the spike protein would affect its site-specific glycosylation pattern. The Alpha variant derived from the D614G lineage is distinguished from others by having deletion mutations located right within an immunogenic supersite of the spike N-terminal domain that make it refractory to most neutralizing antibodies directed against this domain. Despite maintaining an overall similar structural conformation, our mass spectrometry-based site-specific glycosylation analyses of similarly produced spike proteins with and without the D614G and Alpha variant mutations reveal a significant shift in the processing state of N-glycans on one specific N-terminal domain site. Its conversion to a higher proportion of complex type structures is indicative of altered spatial accessibility attributable to mutations specific to the Alpha variant that may impact its transmissibility. This and other more subtle changes in glycosylation features detected at other sites provide crucial missing information otherwise not apparent in the available cryogenic electron microscopy-derived structures of the spike protein variants.

## Introduction

In slightly more than a year’s span since the pandemic outbreak of COVID-19, the SARS-CoV-2 spike glycoprotein (hereafter S protein) has become one of the most intensely studied glycoproteins in the shortest period of time. All possible aspects have been investigated, from its three-dimensional (3D) structure to site-specific glycosylation pattern, complemented by numerous host receptor (human angiotensin converting enzyme 2; hACE2) and neutralizing antibody binding data^1, 2^. The initial effort was to identify any structural difference between the trimeric S protein of SARS-CoV-2 and that of its closest relative, SARS-CoV-1, as well as other related coronaviruses (CoVs) infectious to human. More infectious genetic variants of SARS-CoV-2 have since emerged, as the virus continues to spread and accumulates mutations^3, 4^. The D614G mutation in the S protein was among the first identified in the early phase of the pandemic that became prevalent globally with increased infectivity^5–7^. Starting late 2020, rapid evolution led to simultaneous appearance of several variants of concern^8^, including the B.1.1.7 variant first appeared in the UK, now labeled as the Alpha variant^9^, which was found to be more transmissible and likely also caused more severe illness^10, 11^. In fact, each emerging variant imposes considerable uncertainty factors on the effectiveness of currently available vaccines and neutralizing antibodies^3, 4, 12^, all of which were initially developed based on the original non-mutated spike protein sequence.

Currently available glycosylation data for the intact S protein are mostly derived from LC-MS/MS-based analysis of the recombinant trimeric samples likewise constructed based on the original non-mutated ectodomain sequence and produced in HEK293^13–17^ or insect cells^18–20^. Since glycosylation is both host and protein conformation-dependent, it is anticipated that the glycosylation pattern of HEK293-derived intact trimeric S proteins would better resemble that of the actual virus infecting and propagating through human host cells, particularly at the level of the relative amount of oligomannose versus hybrid/complex type at each of the known 22 N-glycosylation sites. Beyond this simplistic classification, further enumerating the exact terminal structural units carried on the complex glycans may not be relevant since it would be HEK293-specific and not reflecting that of the infected cells and infecting virus *per se*. HEK293, for example, is known to produce significant amount of terminal ±sulfated GalNAcβ1-4GlcNAc (LacdiNAc), with little or no Lewis type structures^14, 21^, which are not universal features of all human cells. Similarly, glycosylation analyses of S proteins produced in insect cells, or the S1 and S2 subunits^17, 20, 22^ independently expressed out of their trimeric assembly context, only matter in so far as considering their use as antigens for diagnostics and vaccine candidates. A serological assay^23^ has demonstrated that the S protein produced in HEK293 cell displays a significantly higher reactivity in COVID-19 plasma/serum detection compared to that produced in insect cells, despite the two sharing an identical protein sequence, thus underscoring the importance of the glycosylation patterns to host immunity and potentially vaccine efficacy.

Since the pandemic outbreak, many laboratories, including ours, have been engaged in determining the atomic structures of the aforementioned trimeric S proteins produced in HEK293, along with rapid site-specific glycosylation mapping, not least to ensure batch-to-batch consistency of antigen production for various downstream assays, diagnostics and vaccine development. Typically, to increase stability and improve yield, the soluble form of intact S protein trimer was engineered to have the putative furin cleavage site abolished together with a tandem proline replacement in the S2 subunit to lock the S protein in a prefusion conformation (known as the 2P mutation), the transmembrane helix was replaced by a T4 fibritin foldon trimerization motif followed by different affinity tags for purification. More recent work has demonstrated that the degree of downstream processing of glycosylation at each site closely resembles that derived from virion particle, when side-by-side analysis was performed^17^. This lends credence to analysis of the engineered recombinant trimer as surrogates of the trimeric S protein on true virion. Any significant departure from the established oligomannose versus hybrid/complex type N-glycans distribution on each site may be interpreted as altered local conformation and spatial flexibility that would affect the accessibility of various glycosyltransferases. This is particularly relevant as mutated variants are evolving rapidly. LC-MS/MS-based glycosylation analysis is far more sensitive, rapid, accessible, and less costly than producing sufficiently good quality and quantity of recombinant proteins for cryo-electron microscopy (cryo-EM) structure determinations. Moreover, there are already several reports showing that the glycan coats may act on the conformational plasticity of the S protein beyond a simple shielding effect^24, 25^. Others have shown that glycans, particularly those on the N-terminal domain (NTD), would potentiate binding to auxiliary lectin receptors, thus enhancing infectivity^26, 27^. While none of the accumulated mutations in the variants of concern directly abolish any of the conserved 22 N-glycosylation sites, it is not known if their respective glycosylation pattern would be significantly altered by underlying conformational changes, which, in turn, may impact on the viral transmissibility, pathogenic severity of COVID-19, and the preventive solutions being deployed worldwide.

To allow a reliable site-specific glycosylation mapping comparison across different S protein constructs and mutated sequences, a standardized and reliable analytical platform is required to discern any real shift in the glycosylation pattern of the variants from random noise intrinsic to the very heterogeneous nature of protein glycosylation. Drawing from the many reported analyses and our own, we have carefully evaluated the S protein glycopeptide data acquisition and analysis settings. We then adopted a sample and data processing workflow that places a higher premium on throughput and objective consistency over comprehensive coverage in both qualitative identification and quantitative mapping. Biological triplicate analyses starting from producing each recombinant spike protein were undertaken, and glycosylation variations were faithfully recorded before averaging and normalization. We further dealt with problematic technical issues, including balancing the false positives and negatives and identifying the minor population of O-glycopeptides amidst the overwhelming pool of N-glycopeptides and peptides without additional enrichment or fractionation.

We found that the mutations of the D614G and Alpha variants did not impact significantly on the gross site-specific glycosylation pattern, consistent with no major change in the 3D conformation of all variants analyzed to date by cryo-EM. Focusing on the few key N-glycosylation sites in proximity with the pocket formed by the receptor-binding domain (RBD) and the NTD of one protomer and the neighbouring RBD from another protomer, we nevertheless detected a significant shift in the relative amount of less processed oligomannose state versus more processed complex type structures at a few sites, indicative of a slight but significant alteration in the local spatial compactness and accessibility. Of high relevance is the considerable shift to more processed N-glycans at site N122 in the Alpha variant, which is directly affected by the nearby Y144 deletion (Δ144), as evident from the cryo-EM density map. Moreover, preserving the original furin processing ^682^RRAR^685^ sequence indeed brought instability and lower yield but curiously effected a change in the predominant O-glycosylation forms of one site without significantly altering the overall N-glycosylation pattern. We conclude that minor changes brought by deletion and point mutations in local conformation within a largely stable overall footprint would considerably affect the downstream processing efficiency of the attached glycans. Since each of the glycans covers a not-so-insignificant surface area and may effectively mask or distort a conformational epitope, as well as engaging additional host receptors, its impact on the efficacy of neutralizing antibodies and the acquired cell-mediated immunity upon vaccination cannot be under-estimated.

## Material and Methods

### Plasmid construction and purification of SARS-CoV-2 Spike

The codon-optimized nucleotide sequence of full-length SARS-CoV-2 S protein and that of the Alpha variant were kindly provided by Dr. Che Alex Ma (Genomics Research Center, Academia Sinica) and Dr. Mi-Hua Tao (Institute of Biomedical Sciences, Academia Sinica), respectively. The DNA sequence corresponding to residues 1-1208 of the S protein was subcloned from the full-length S sequence, appended with the T4 fibritin foldon sequence at the C-terminus, followed by a c-Myc sequence and a hexahistidine (His_6_) tag, and inserted into the mammalian expression vector pcDNA3.4-TOPO (Invitrogen, USA). The D614G mutant was generated by site-direction mutagenesis with specific primers as described in our previous study^28^. Two variants were subsequently generated by introducing a tandem proline stabilization mutation (2P, ^986^KV^987→986^PP^987^), with and without furin cleavage site mutation (fm, ^682^RRAR^685→682^GSAG^685^), designated as S-fm2P and S-2P, respectively. The same construct design of S-fm2P was used for the D614G and Alpha variants (hereafter designated as S-D614G and S-Alpha, respectively).

The expression vectors encoding for all S proteins were transiently transfected into HEK293 Freestyle (HEK293F) cells with polyethylenimine (PEI, linear, 25 kDa, Polysciences, USA) at a ratio of DNA: PEI = 1:2. The transfected cells were incubated at 37°C, with 8% CO_2_ for six days. The cells were pelleted by centrifugation at 4000 rpm for 30 min, and the supernatant containing the overexpressed S proteins was collected and filtered through a 0.22 μm cutoff membrane (Satorius, Germany). The supernatant was incubated with HisPur Cobalt Resin (Thermo Fisher Scientific, USA) in 50 mM Tris-HCl (pH 8.0), 300 mM NaCl, 5 mM imidazole, and 0.02% NaN_3_ at 4 °C overnight. The resin was sedimented by gravity in an open column (Glass Econo-Column® Chromatography Column, Bio-Rad, USA), washed with 50 mM Tris-HCl (pH 8.0), 300 mM NaCl, 10 mM imidazole, and the target protein was eluted by 50 mM Tris-HCl (pH 8.0), 150 mM NaCl, 150 mM imidazole. The eluent was concentrated and further purified by using a size-exclusion chromatography (SEC) column (Superose 6 increase 10/300 GL, GE Healthcare, USA) in 50 mM Tris-HCl (pH 8.0), 150 mM NaCl, 0.02% NaN_3_. The purity of the samples was confirmed by SDS-PAGE. The protein concentrations were determined by using the UV absorbance at 280 nm using a UV-Vis spectrometer (Nano-photometer N60, IMPLEN, Germany).

### SEC-MALS

The integrity of the trimeric assembly of the S-fm2P and the degree of glycosylation was confirmed by SEC coupled with multiangle light scattering (SEC-MALS) as described previously^29^. The purified S protein was separated by an SEC column (BioSEC-3, Agilent, USA) connected to an HPLC system (Analytical HPLC 1260 LC system, Agilent, USA), and the corresponding static light scattering signals were detected by a Wyatt Dawn Heleos II multiangle light scattering detector (Wyatt Technology, USA). To dissect the contributions of amino acids and glycans to the overall molecular weight of the SEC elution peak, the refractive index increments (dn/dc) of protein and protein conjugate (glycan moiety) were defined as 0.185 and 0.140 mL/g, respectively. The buffer viscosity (η) was set to 0.8945 cP at 25 °C based on the theoretical estimate using SEDNTERP. Those parameters were applied in ASTRA 6.0 software (Wyatt Technology, USA).

### In-solution proteolytic digestion of the S proteins

Proteins were reduced with 10 mM dithiothreitol at 37 °C for 1 hr, then alkylated with 50 mM iodoacetamide in 25 mM ammonium bicarbonate and 4 M urea for 1 hr in the dark at room temperature. After that, dithiothreitol was added to a final concentration of 50 mM, then buffer-exchanged to 25 mM ammonium bicarbonate buffer using Amicon Ultra-0.5, 10 ka (Merck Millipore, Ireland) and treated overnight with sequencing grade trypsin and chymotrypsin (Promega, USA) simultaneously at an enzyme-to-substrate ratio of 1:30 at 37 °C. The digested products were then diluted with formic acid to a final concentration of 0.1% for further experiments.

### Glycopeptides analysis by liquid chromatography-mass spectrometry

The digested peptides were cleaned up using ZipTip C_18_ (Merck Millipore, Ireland), dried down, and redissolved in 0.1% formic acid (Solvent A). Data were acquired on Orbitrap Fusion Lumos Tribrid mass spectrometer (Thermo Fisher Scientific, USA) fitted with an Easy-nLC 1200 system (Thermo Fisher Scientific, USA). For each LC-MS/MS analysis, an equivalent of 1 μg glycoprotein digest was loaded onto an Acclaim PepMap RSLC C_18_ column (Thermo Fisher Scientific, Lithuania) and separated at a flow rate of 300nL/min using a gradient of 5% to 40% solvent B (80% acetonitrile with 0.1% formic acid) in 200 min. The mass spectrometer was operated in the data-dependent mode. Briefly, survey full scan MS spectra were acquired in the Orbitrap from 400 to 1800 m/z at a mass resolution of 120,000. The highest charge state ions within charge state 2 to 6 were sequentially isolated for MS^2^ analysis using the following settings: HCD MS^2^ with AGC target at 5×10^4^, isolation window 2, orbitrap resolution 30000, step collision energy (%): 25, 28, 32. HCD-pd-EThcD triggered by at least detection of one of the product ions at *m/z* 138.0545, 204.0867, 274.0926, or 366.1396 within the top 20 ions. For EThcD, the calibrated charge-dependent parameter was used, the supplemental activation collision energy was 15 %, and orbitrap resolution 60000. For each of the SARS-CoV-2 S protein variants, three biological replicates were prepared and analyzed by LC-MS.

### Glycopeptide Identification and quantification

The MS and MS^2^ data were first evaluated by Preview (Protein Metrics Inc., USA) to determine the optimum search parameters for Byonic (Protein Metrics Inc., USA). The HCD and EThcD MS^2^ data were then processed by Byonic (v 3.10.10 for S-2P/fm2P/D614G O-glycopeptide identification, and v 3.11.3 for S-Alpha O-glycopeptide and all N-glycopeptide identification) using the following general parameters: search against the SARS-CoV-2 spike protein sequence with semi specific cleavages at F, Y, W, L, K, R residues, allowing up to 2 missed cleavages, with the precursor ion mass tolerance set at 4 ppm and the fragment ion mass tolerance at 10 ppm. Fixed modification included cysteine carbamidomethylation (+57.0215 Da, at C), whereas variable common modifications considered were carbamidomethylation (+57.0215 Da, at H, K), dithiothreitol (+151.9966 Da, at C), oxidation (+15.9949 Da, at M), deamidation (+0.9840 Da, at N), carbamylation (+43.0058 Da, at N-terminus, K, R), and variable rare modifications considered were Gln to pyro-Glu (−17.0265 Da, at N-terminal Q), Glu to pyro-Glu (−18.0106 Da, at N-terminal E). The N-glycopeptide and O-glycopeptide identification were processed by Byonic separately. For N-glycopeptide identification, the built-in N-glycan library of “132 human” was used as a base but modified by i) removing N-glycans smaller than trimannosyl core structure (HexNAc_2_Hex_3_), and ii) adding in 8 N-glycans not listed in the library and selected N-glycan compositions carrying an extra sulfate (see Supplemental Table S1 for the exact N-glycan library used in this work). For O-glycopeptide identification, the built-in O-glycan library of “O-glycan 9 common” was used to identify O-glycopeptides. The criteria used in additional manual filtering of positive matches were PEP2D<0.001, score >200, and retaining only those identified by such criteria in at least 3 out of the 12 datasets analyzed. The filtered Byonic identification results were output into an Excel file (Table S2) along with curated non-redundant lists of identified glycan compositions and peptide backbones to be used as one of the filtering criteria for subsequent quantification output.

The unfiltered Byonic search results were fed to the Byologic module of the Byos suite (v3.11, Protein Metrics Inc., USA) for quantification purpose based on peak areas of extracted ion chromatograms at 5 ppm accuracy. In cases when the same glycan composition of a particular N-glycosylation site occurs on more than one unique peptide backbones due to different peptide modification and/or mis-cleavage, the peak areas for those glycopeptides starting with the same N-terminal amino acids were summed and treated as one quantified entry for that site. Only those quantifiable peaks identified at more relaxed criteria (PEP2D<0.01) but carrying either the glycan and/or peptide backbone identified initially by PEP2D<0.001 and score>200 were exported into an Excel file (Table S3), from which the bar chart and heat maps were generated. For these final outputs, peak areas of all glycopeptides carrying the same glycan composition on a particular site but different peptide backbones were summed as one unique quantified entry. To generate the pie chart for each site, values from three biological triplicates of the same CoV-2 variant were averaged. The O-glycopeptide quantifications (reported in Table S4) were manually performed using the peak areas of the extracted ion chromatograms within 5 ppm accuracy by Xcalibur (Thermo Fisher Scientific, USA), with spurious or non-verifiable peaks excluded.

## Results and Discussion

### Datasets generated and glycopeptide identification criteria

Site-specific glycosylation mapping was undertaken for each of the recombinant trimeric S proteins produced in HEK293F cells, purified from the culture supernatant, checked for the integrity of trimeric assembly by SEC-MALS in the case of S-fm2P (SEC elution profiles for other variants), and used for cryo-EM-based structural studies. Three biological replicates were prepared for each of the S-2P, S-fm2P, S-D614G and S-Alpha variants (Fig. 1A), yielding a total of twelve S protein samples subjected to an analytical workflow (Fig. 1B) optimized for rapid and standardized quantitative assessment of potential variations in site-specific glycosylation. Our strategy did not aim for comprehensive coverage of all 22 known N-glycosylation sites, nor specifically gear towards uncovering as many low occupancy O-glycosylation sites. Instead, a single simultaneous trypsin and chymotrypsin digestion was applied, which would reproducibly yield good quality site-specific N-glycosylation information for 19 sites, including the few critical ones for the NTD and RBD. Data visualization for the N-glycosylation pattern is provided in a few different formats. Firstly, the complete qualitative identification results as processed by Byonic search and filtered by PEP2D<0.001 and score>200, and the ensuing Byologic-quantifiable glycopeptides filtered by the unique peptide backbone and glycan composition derived from the higher stringency identification results, were reported as Tables S2 and S3, respectively, in Excel sheets format. All twelve LC-MS/MS raw datasets have also been deposited to MassIVE site. Secondly, the quantified glycopeptides normalized as percentage total of all glycan composition identified for each site were visualized as a comprehensive heatmap and presented in Fig. S1. This is further supplemented by individual zoomed-in heat maps for sites N74, N122, and N282 (Fig. S2), after removing glycopeptides not found on these particular sites.

**Fig 1.**
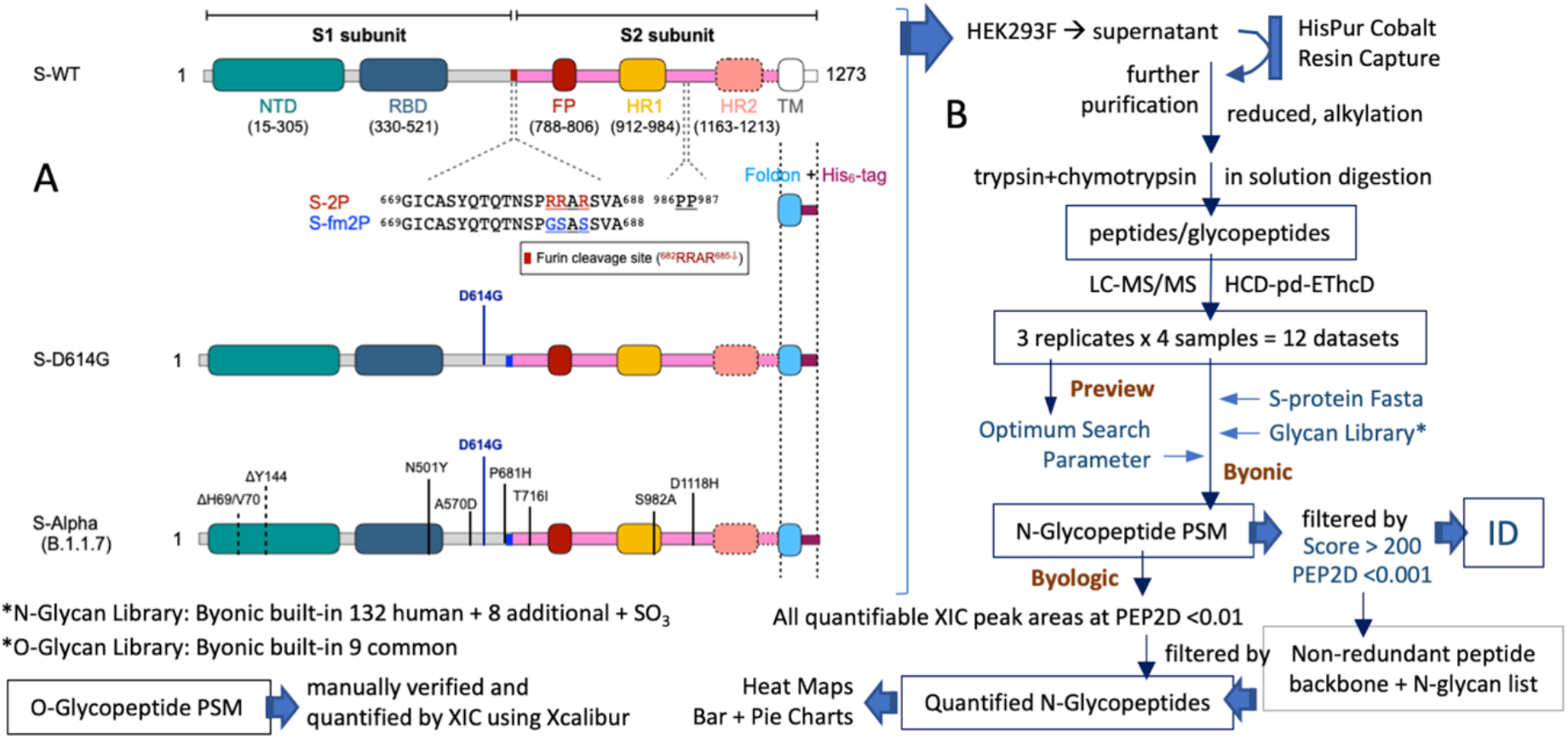
The construct designs for the recombinant trimeric spike proteins and the analytical workflow used to map their site-specific glycosylation profiles. **(A)** The trimeric prefusion S proteins analyzed in parallel for direct comparison were all stabilized by 2P mutations and RRAR to GSAS substitution at the furin cleavage site, with a foldon trimerization motif added for the wildtype (fm2P), D614G and Alpha (B1.1.7) variants, but a stabilized wild type S protein (2P) retaining the furin cleavage site was also included. Despite the deletions in the S-Alpha, the residue numbers of the remaining protein sequence and the N-glycosylation sites follows that of S-WT and the other variants without deletions. **(B)** The recombinant S proteins were purified from the culture supernatant and digested simultaneously by trypsin and chymotrypsin for LC-MS/MS analysis as indicated. The acquired datasets were processed by Byonic and the Byologic module under Byos® for glycopeptide identification and quantification, respectively. N- and O-glycopeptides were searched and analyzed separately for better results.

Thirdly, normalized bar charts for the relative amount of major glycopeptide types identified for select NTD sites on each of the four S protein variant triplicates are presented to enable rapid visualization of how the site-specific glycosylation pattern, including the degree of sialylation, fucosylation and sulfation may differ among different S protein variants and among the sample triplicates (Fig. 2A). Fourthly, a simplified pie chart version grouping the glycopeptides into different oligomannose types (Man5-Man_9_HexNAc_2_) and complex/hybrid types containing 3-7 HexNAc (disregarding the number of Hex, Fuc, and NeuAc) for 15 sites are given for comparison across the three furin site mutated S protein variants (Fig. 2C). Unlike the bar charts that faithfully show the batch-to-batch variation, these pie charts utilized averaged data from the triplicates for each protein sample, which is more representative but at the expense of degenerated information on glycosylation details and variations. Finally, the most abundant N-glycan composition for each identified site of the Alpha variant was presented in cartoon drawings and localized onto an S-protein structure rendered from the cryo-EM data to indicate how these are spatially distributed (Fig. 2B).

**Fig 2.**
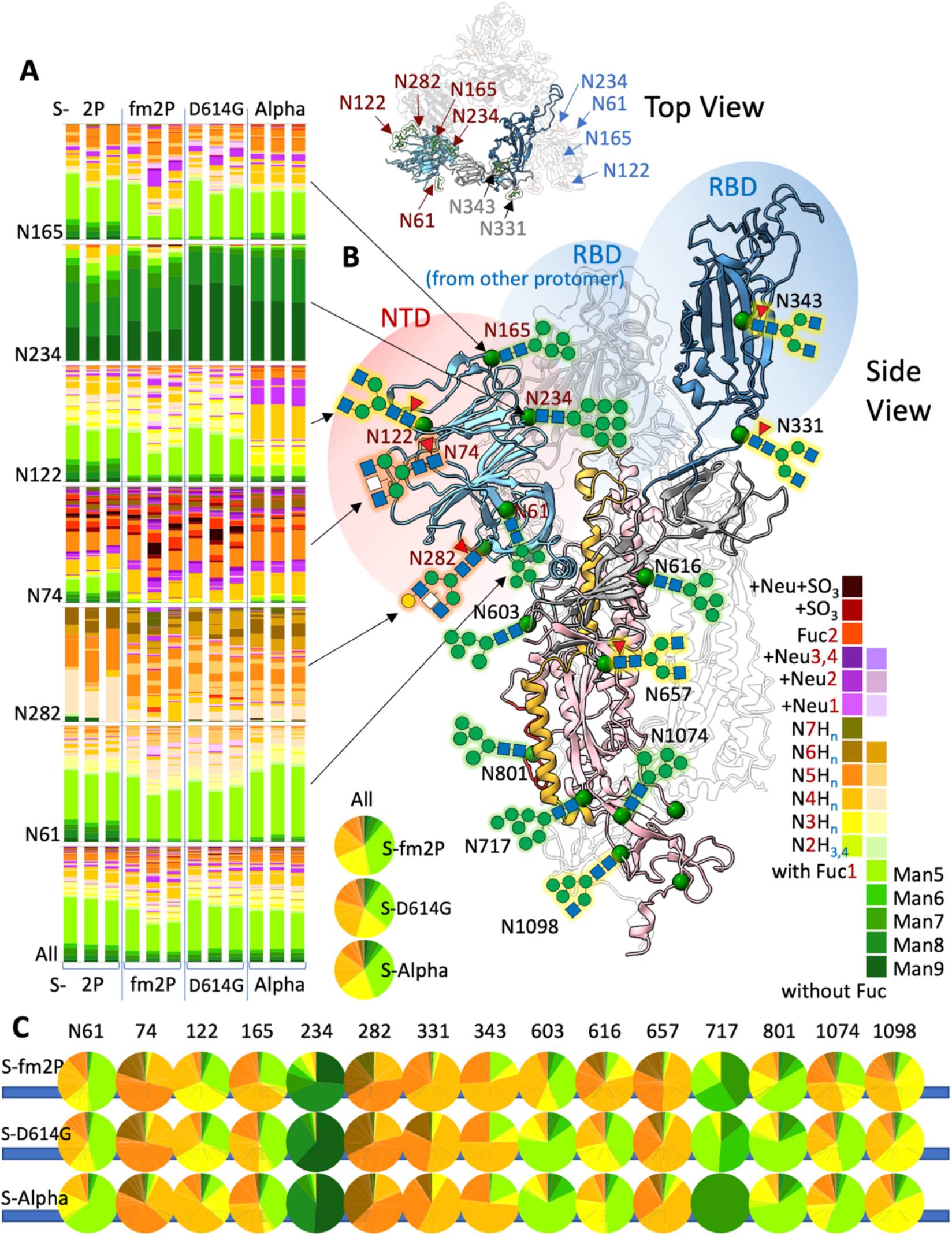
Overall site-specific glycosylation pattern of the recombinant trimeric spike proteins derived from the original SARS-CoV-2 strain and its mutated D614G and Alpha variants. **(A)** Full glycosylation heterogeneity for the few NTD sites as revealed by triplicate analyses of the 4 samples including the S-2P sample retaining the RRAR furin cleavage site. Their respective locations on the overall 3D structure of the spike protein were indicated. **(B)** Side view and top view of the trimeric S proteins based on our cryo-EM structure of S-D614G (PDB ID 7EAZ), highlighting a single protomer with another protomer (faded) in the background. The most abundant N-glycan identified for each N-glycosylation site of S-Alpha is depicted by cartoon drawing adopting the recommended Symbol Nomenclature for Glycan (SNFG)^41^. White square indicates a HexNAc, which can either be GalNAc if the structure carries terminal LacdiNAc (GalNAcβ1-4GlcNAc-) unit or GlcNAc if it is structure with terminal non-galactosylated GlcNAc and/or bisecting GlcNAc. **(C)** Data from triplicate analyses were averaged and degenerated into Man_3-9_GlcNAc_2_ and HexNAc_4-7_X (X can be any number of Hex, Fuc, NeuAc and sulfate) to be visualized as pie charts for 15 out of 17 sites with quantifiable N-glycopeptides. The color code used for both (A) and (C) is shown as strips of color chart. In (A), any structure with sulfate (SO_3_), NeuAc (Neu), or more than one Fuc will be color coded as such, disregarding the number of HexNAc (N) and Hex (H). The glycan compositions are arranged by increasing number of HexNAc, from bottom to up in the bar charts. Full representation in heat maps can be found in Figs. S1 and S2.

As already discussed by others, there are many sources that would contribute to discrepancies among the reported glycosylation patterns, including the exact recombinant protein constructs, host cell culture conditions, protein purification and digestion methods, the LC-MS/MS data acquisition, processing and analysis parameters, as well as the criteria used to accept or filter the computational search results. To strike a fine balance between data processing throughput and accuracy, as well as objective consistency, N-glycopeptides identification by Byonic for S-fm2P and S-2P were first scrutinized for the numbers of peptide spectrum matches (PSM) containing at least three peptide fragment ions at PEP2D<0.001 coupled with different score cut-off (Fig. S3), which led to adopting the additional filtering criteria of score>200 to be applied across all datasets. The glycan compositions thus identified on each site were then manually checked for any inconsistency against the well-known glycomic repertoire of HEK293^21^. During this process, a common source of error in misassigning the mass difference between HexNAc and Fuc to carbamidomethylation on Lys by Byonic due to the absence of the correct glycan composition in the 132 N-glycan library used was identified (Fig. S4) and rectified by adding in 8 additional entries (Table S1), for subsequent reprocessing and processing of all datasets. We further confirmed, by the presence of diagnostic oxonium ions at *m/z* 407.166, that terminal HexNAc_2_ unit attributable to LacdiNAc is a common feature, along with ions that respectively verify the occurrence of sulfated HexNAc and fucosylated HexNAc_2_ particularly on, but not restricted to, glycopeptides from site N74 (Fig. S5). This glycosylation characteristic of HEK293F is reflected by several identified glycan compositions with Fuc2 and even Fuc3, or sulfate, with and without additional sialylation, as highlighted in the color code used (Figs. 2A, S1, S2).

### Consistent features among the variations

Due to substantial inherent variations in sample and data processing among different laboratories, detailed comparison of identified site-specific N-glycopeptides down to individual glycoforms level is not possible and largely non-meaningful. On the other hand, a few emerging characteristics homing in on the overall degree of glycan processing on specific sites as inferred from the relative amount of oligomannose versus hybrid/complex type structures appear to be very consistently reproduced by each reported analysis to date. Allen *et al.*^16^ has applied the same data processing method to analyze the available datasets from 5 different sources of prefusion state stabilized recombinant trimeric spike proteins (Amsterdam Medical Centre, Harvard Medical School, Switzerland, The Wellcome centre for Human genetics, and University of Southampton/University of Texas at Austin), very similar to our wildtype S-fm2P used here. They further calculated the average oligomannose/hybrid versus complex type glycan compositions carried on each site from these 5 samples. In addition, Wang *et al.*^15^ analyzed the same recombinant trimeric S protein (University of Texas at Austin) first examined by Watanabe *et al.*^13^. More recently, Brun *et al.*^17^ analyzed a different stabilized source of recombinant trimeric S protein (Ichan School of Medicine at Mount Sinai, New York). All these cited works similarly used Byonic/Byologic for quantitative mapping of intact trimeric S protein ectodomain produced in mammalian cells (HEK293, with the exception of the Swiss sample produced in CHO cells). Collectively, these provided a good reference set of data to be compared against our own Academia Sinica produced sample (Fig. 3). Another report by Zhao *et al.*^14^ used spectral count (PSM) for quantification, which makes direct comparison difficult, although the overall picture concluded is similar. Its source of the recombinant S protein (Harvard Medical School) was, however, already included in the recent work by Allen *et al.*^16^. Other reports on S1/S2 protomer or subunits^17, 20, 22^, RBD domain^20, 30–32^, or intact S protein produced in insect cells^18–20^ are not considered here since these carried distinctively different glycosylation pattern.

**Fig 3.**
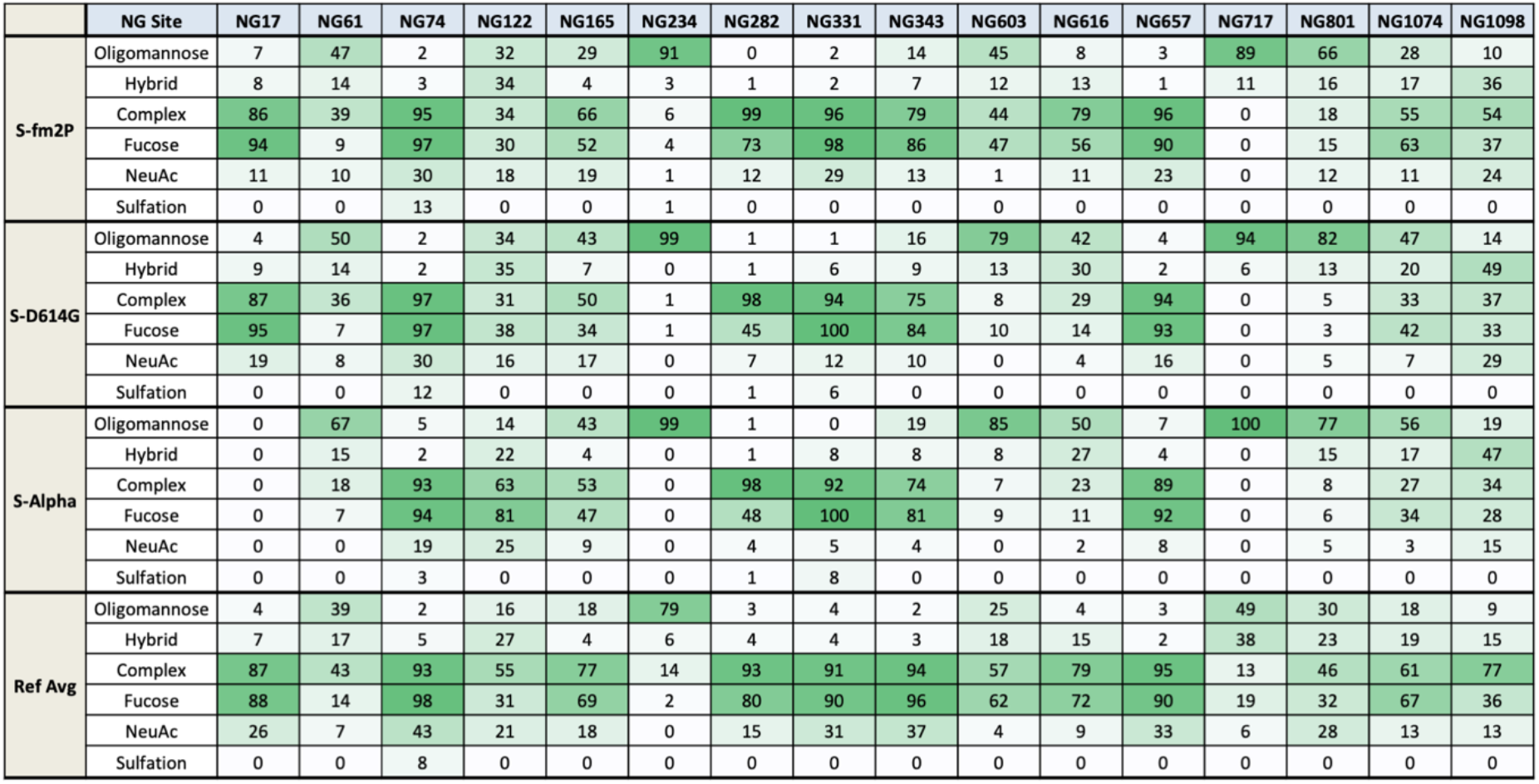
The relative abundance of major classes of site-specific N-glycopeptides identified on the stabilized trimeric S-protein variants. The site-specific N-glycopeptides were classified according to their assigned glycan compositions, with Man_5-9_GlcNAc_2_ defined as oligomannose type, HexNAc_3_-X as hybrid type, HexNAc_4-7_-X as complex type, with X can be any number of Hex, Fuc, NeuAc and sulfate. The definition of HexNAc_3_-containing N-glycan as hybrid type does not take into considerations that some HexNAc_4-5_-containing N-glycans can also be hybrid instead of complex type, which cannot be readily resolved by current MS^2^ data on the glycopeptides. The numbers in each cell refer to the % total after summing the quantified peak areas for the respective classes of identified N-glycopeptides, using the average value from triplicate analysis (see Table S3). Fucose, NeuAc and sulfate refer to N-glycopeptides identified as carrying one or more of these substituents and calculated as % total over all quantified N-glycopeptides for that particular site. The color intensity reflects the relative abundance. Data for S-fm2P, S-D614G and S-Alpha were generated in this work. These are compared against the average value (Ref Avg) derived by Allen *et al.*^16^ from datasets of five different sources of similarly prefusion state stabilized recombinant trimeric spike proteins analyzed by others. For direct comparison, the values for Hybrid, Fhybrid, HexNAc_3_x and HexNAc_3_Fx in their chart were summed together as Hybrid in this Table and the % total values for oligomannose, hybrid and complex were recalculated accordingly.

### All complex types are not made equal

Focusing on the NTD, the most consistent glycosylation pattern is the one at N234, which is the only site that is always >80% unprocessed at Man8 and Man9-oligomannose state. Like the Harvard and Swiss samples, our own S-fm2P sample yielded a small % of complex type glycan at this site, whereas samples from other sources, as well as our D614G and Alpha variants, are approaching 100% oligomannose, which may reflect a slight difference in the spatial accessibility afforded by the different trimeric assembly. On the other extreme, glycans at N74 and N282 are consistently processed almost completely to complex type structures for all samples. In fact, these two NTD sites, which are situated away from interacting directly with the RBD from other protomers, carry the most varied glycosylation heterogeneity that differs significantly between the two sites (Fig. S2). N74 is not only more highly sialylated but also is the site carrying the highest amount of sulfation and additional Fuc, both due to the prominence of LacdiNAc (Fig. S5). In contrast, a significant proportion of glycans at N282 does not contain Fuc, and most are not sialylated. The average values provided by Allen *et al.* are 98% fucosylated and 43% sialylated for N74 versus 80% and 15%, respectively, for N282. Our own S-fm2P sample yielded even more contrasted values (Fig. 3). This feature is largely maintained by the D614G and Alpha variants although there is a significant decrease in the degree of fucosylation at N282 going from the wild type (both our own and the ref average at 73-80%) to the variants (45-48%). Moreover, the exact distribution and relative amount of each distinct complex type glycan also varied slightly from batch to batch and more pronouncedly from one to another mutated variants. The immunobiological consequence is not possible to determine since the actual glycosylation structures on the infectious virion will be dependent on the infected host cell types. Nevertheless, it points towards the possibility of carrying different glycotopes that can be recognized differently by the glycan-binding proteins on myriad immune cells in infected individuals carrying natural variations in glycomic expression, which would in turn contribute to different immune response and pathological severity.

### Different shades of glycan processing

Sites N61, N122, and N165 typify another category of N-glycosylation sites, which carry a mixture of oligomannose/hybrid and complex type structures at variable amount. The average values provided by Allen *et al.* for oligomannose versus hybrid/complex type structures at these three sites for the 5 recombinant protein samples are 39%:60%, 16%:82% and 18%:81%, respectively^16^. However, there are significant inter-sample variations, whereby oligomannose structures at site N61 can be as high as 70-80% (in Wellcome Center, Southampton/Texas, and Mount Sinai samples) and the complex type structures at site N165 can be as high as approaching 100% (in Swiss and Southampton/Texas samples). Nonetheless, in most cases, including our analysis, the single most abundant structure at N165 is Man_5_GlcNAc_2_. This is also the structure used to model how it is modulating the conformational dynamics of the RBD, along with the Man8 or Man9-oligomannose structure at N234^24^. We observed only a slight increase in oligomannose structures in the overall similar glycosylation pattern of N165 upon D614G and further mutations in the Alpha variant, albeit not without some variations in the actual complex type structural heterogeneity. Similarly, the single most abundant structure at site N61 in most samples is Man_5_GlcNAc_2_. Relative to N165, the degree of fucosylation and sialylation at N61 is much lower, and there appears to be also a slight shift towards more Man_5_GlcNAc_2_ and even less sialylation in the Alpha variant.

The glycosylation pattern at N122 is more similar to N165 than N61 in terms of the extent of processing to complex type structure and degree of sialylation but is significantly less core fucosylated, according to the average value by Allen *et al.* and also reflected in our own analysis. Interestingly, while single mutation at D614G retains much of the same glycosylation pattern, a significant shift is noted in the Alpha variant more than in any other sites. The overall shift (Fig. 4C) is consistent with its gaining a more downstream processing status from Man5 to more highly fucosylated and sialylated complex type structures. On average, the heterogenous glycans at N122 would thus be expected to occupy a bulkier spatial volume in the Alpha variant relative to D614G or the original non-mutated strain. It is perhaps no coincidence that N122 is located nearby a disordered loop (N3, residues 141-156)^33^ that harbors the deleted Y144 in the Alpha variant and is no longer apparent in our reconstructed cryo-EM map (Fig. 4A). The distinct orientation of the N-glycan stubs at N122 further indicates some subtle local conformational changes not apparent from comparing the overall end state structures of the S protein variants resolved by cryo-EM single particle reconstruction. Yet, the extent of site-specific glycan processing as the trimeric spike protein transits through the Golgi apparatus, including the antennary branching, core and additional peripheral fucosylation, sialylation, and sulfation, is extremely sensitive to the slightest change in the local conformational dynamics brought about by the primary sequence mutations. Some sites exemplified by N122 would become more accessible to the Golgi-resident glycosyltransferases, while other sites such as N61 would become less processed. It is not a one-way loosening up of the entire compact structure but rather subtly affected in either way while maintaining the overall structural similarity. These findings exemplify the sensitivity of complementary MS analysis in probing conformational changes during glycan processing that is otherwise inaccessible to other structural biology tools, including cryo-EM.

**Fig 4.**
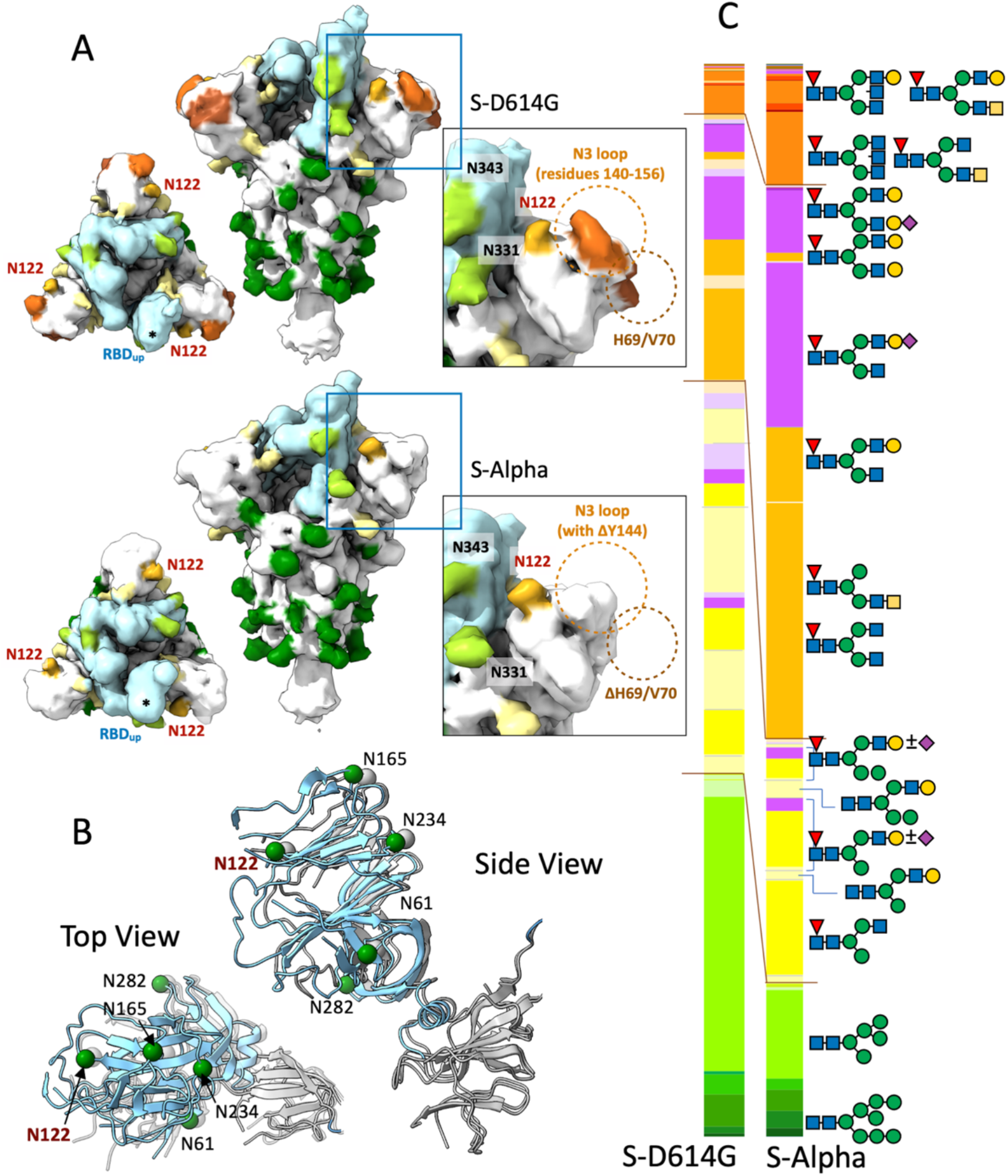
N-glycosylation at site N122 and its location at an exposed region of the N-terminal domain of the trimeric S protein variants. **(A)** Orthogonal views of the low-pass filtered cryo-EM maps of S-D614G and S-Alpha with the visibly resolved stems of N-glycans (first two GlcNAc moieties extending from the sidechain of Asn) at specific regions highlighted in different colors. The five N-glycans within the NTD are colored in light yellow, with the exception for N122, which is colored in gold. The N-glycans at N331 and N343 within the RBD are colored in lime. The expanded views of the NTD highlight the differences in the EM maps between the D614G and Alpha variants due to Δ69-70 and Δ144 in the latter. **(B)** Orthogonal views of the superposition of the atomic models of the D614G and Alpha variants, which are shown in gray (PDB ID 7EAZ) and cyan (PDB ID 7EDF), respectively. The positions of the backbone Cα atoms of the N-glycosylated asparagine residues are shown in green or gray spheres with their residue identities indicated. **(C)** The glycosylation bar charts for N122 on the S proteins of the D614G and Alpha variants, averaging data from triplicate analysis. The most likely N-glycan structures corresponding to select few of the identified glycosyl compositions are annotated as cartoon drawings using the SNFG standard. These structures were not further verified experimentally. The color code used is the same as the one shown in Fig 2.

### All shifts are towards less processed states except N122

The N-glycans at N331 and N343 located in the RBD have consistently been reported as highly processed, implying a relatively exposed accessibility. In our hands, N343 does have a few oligomannose structures (~15%) but, importantly, both sites carry mostly complex type structures and are not appreciably affected by mutations in the D614G and Alpha variants. Outside the RBD, site N657 carries mostly complex type structures, while both N603 and N616 have a mixture of oligomannose and complex type structures with the former consistently less processed. Notably, the value for N603 (and several other sites) as determined from analyses of different S-protein sources varies considerably^16^ and not well represented by the reported average value (Fig. 3), which was skewed by atypically high amount of complex type structures at this site in the Swiss and Harvard samples^16^. Getting into S2, the first few sites from N717, N801 to N1074 spotted a decreasing proportion of oligomannose structures. This trend is equally observed in other reported samples^16^, and upheld in our mutated variants. The remaining N-glycosylation sites towards the C-terminus of S2 are all dominated by complex type structures with few or no oligomannose structures. More importantly, we consistently detected substantial increases in the relative amount of oligomannose structures at N603, N616, and N1074 in the D614G and Alpha variants. The overall picture gleaned from these analyses suggests that S-D614G and S-Alpha are more similar in their N-glycosylation pattern at a majority of sites and, relative to S-fm2P, generally show a shift towards less processed states for sites that originally carry a variable mixture of oligomannose and complex type structures. Only glycosylation at N122 among the sites analyzed bucks this trend and gets more processed into complex types instead. On the other hand, N-glycosylation at the very N-terminal sites N17, N61, N74, and N122, are less affected by the D614G mutation alone but register more pronounced shifts in the S-Alpha variant that carries multiple mutations, including the two deletions in the N-terminal domain.

### Impact of furin site and other mutations on O-glycosylation

It is now known that retaining the RRAR furin cleavage site would adversely affect the yield of the trimeric recombinant S protein produced in HEK293 cells due to potential cleavage into S1 and S2 in Golgi, which may, however be still held together^17, 34^. All cryo-EM and glycosylation analyses of stabilized trimeric S proteins to date were thus performed on S-fm2P derivatives with the site mutated. We indeed found that our S-2P protein could be recovered and purified in a manner similar to other GSAS-stabilized S-fm2P proteins but at a much lower yield and gave bands corresponding to both uncleaved S1/S2 and cleaved S1 and S2 subunits by SDS-PAGE (Fig. S6). Despite using the same total protein amount for digest and LC-MS/MS analysis, the signal intensity was significantly lower than the other three recombinant spike proteins with furin cleavage site mutated. This may have unnecessarily skewed the pattern towards slightly more oligomannose type profiles since glycopeptides with oligomannose structures are more readily identified by the analytical process. The data was therefore not used for direct comparison with those of other S protein variants. Nonetheless, the overall site-specific glycosylation characteristics of S-2P largely recapitulate the overall picture concluded from analyzing the GSAS-mutated S-fm2P, S-D614G and S-Alpha, namely with N-glycans at N234 being almost all retained as oligomannose, N74 and N282 being highly processed into complex types, and N61, N122, and N165 somewhere in between (Fig. 2A). This is consistent with the current understanding that furin-induced cleavage, if occurs, is a rather late Golgi event^17, 35^ after all glycan addition and further processing into complex type structures have largely taken place. It also indicates that having RRAR instead of GSAS did not significantly affect the overall 3D structure of the S protein.

Interestingly, however, we noted that S-2P actually yielded significantly more of mono- and disialylated core 1 and 2 O-glycans at T323 relative to the non-sialylated core 1 and 2 and single GalNAc structures (Fig. 5). This is relevant since sialylated glycopeptides are usually more difficult to identify than non-sialylated ones due to less favorable ionization and fragmentation properties. The fact that a higher proportion of sialylated glycopeptide was identified for a sample of lower amount that yields overall lower signal intensity added confidence that the increase in sialylation at this O-glycosylation site is of real significance. It reflects a relatively late Golgi event that coincides with potential S1/S2 cleavage. Among other O-glycosylation sites, we only detected a low level of O-glycosylation at T678 located within the loop that harbors the furin cleavage site for the GSAS-stabilized S-fm2P but not S-D614G or S-Alpha (Table S4). Incidentally, both variants afforded higher levels of sialylation on their O-glycans at T323, relative to S-fm2P and more similar to S-2P (Fig. 5B). Collectively, these findings indicate that the spatial confinement or accessibility of T323 within the RBD is sensitive to local conformational changes and those occurring elsewhere, including the non-structured S1/S2 loop that may or may not be cleaved before the trimeric S protein exits Golgi.

**Fig 5.**
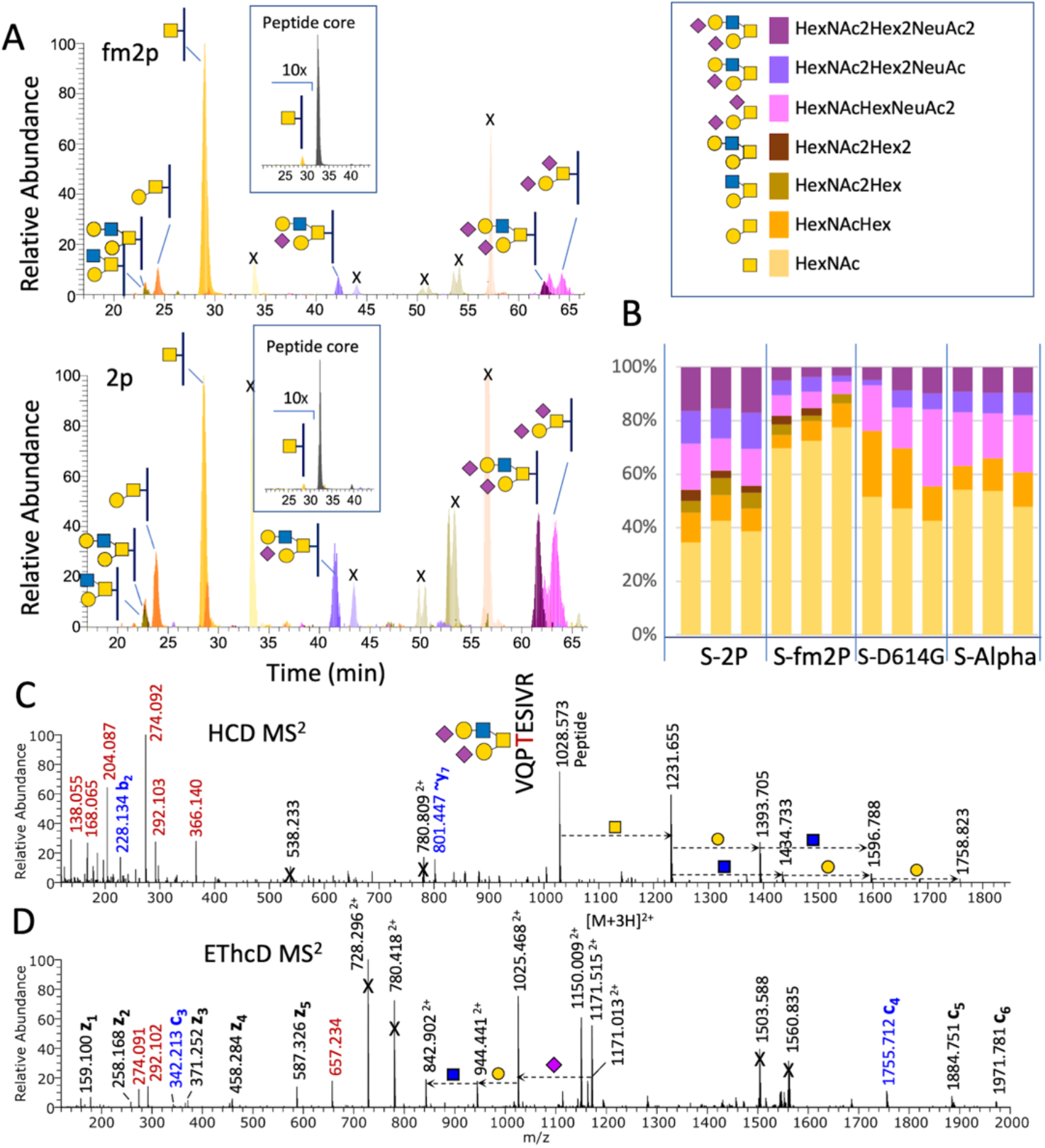
Quantitative analysis of O-glycosylation at T323. **(A)** Extracted ion chromatograms of the peptide ^320^VQP<u>T</u>ESIVR carrying the common cores 1 and 2 O-glycans for S-fm2P (upper panel) and S-2P (lower panel). Each of the putative peaks were manually verified and those not corresponding to the expected glycopeptides are marked with “X”. The overall occupancy is very low and the inset shows the amount of the most abundant glycoforms with single GalNAc relative to the non-glycosylated peptide, after 10x magnification. **(B)** The quantified distribution of various glycoforms for this site for all 12 samples (3 replicates each for the 4 recombinant spike proteins) analyzed, with the color code used for different O-glycan structures shown in the box above. Full data for this and other identified O-glycosylation sites are presented in Table S4. **(C, D)** HCD-MS^2^ and EThcD-MS^2^ data for the disialylated HexNAc_2_Hex_2_-carrying O-glycopeptide from S-2P. The c_3_ and c_4_ ions afforded by EThcD-MS^2^ unambiguously localize the disialylated O-glycan to T323.

### Concluding Perspectives

Despite many reports on the site-specific glycosylation pattern of the SARS-CoV-2 spike proteins in all different states, including S1/S2 protomer, subunit, and stabilized trimer produced either in insect or HEK293 cells, a detailed comparison is difficult. The many sample preparation, analytical data acquisition, and processing variations almost assure a significant level of differences in view of the heterogenous nature of protein glycosylation. Nonetheless, if this heterogeneity is degenerated simply into oligomannose versus hybrid/complex type as a measure of the extent of glycan processing and hence spatial accessibility, a more unifying picture informative of the glycosylation site-specific local conformation of a properly assembled trimeric S protein can be better described.

In that respect, our glycosylation analysis of the S proteins is largely consistent with the currently accepted model^16^. More importantly, this is the first report that attempts to identify any departure from this archetypal view as mutated SARS-CoV-2 variants of concern evolve. It is made possible by having tight quality control over the in-house produced S protein sample source in conjunction with applying the same analytical workflow, data analysis parameters, and objective identification criteria for side-by-side comparison. Based on the overall very similar protein structures determined by companion cryo-EM analysis, large differences in glycosylation pattern are not expected. We nevertheless identified several distinct trends, including a more processed state at N122 in the S-Alpha variant when most other sites shifted instead towards less processed status. Among the sites that are mostly decorated with complex type structures such as N74 and N282, the exact N-glycans carried and their overall degree of sialylation and fucosylation, not only vary between sites but also between the original and mutated variants for the same site. This extends further to O-glycosylation profiles at T323 and T678, although their very low level of occupancy may have much less impact.

The few critical mutations in the RBD of the S-Alpha appear not to have significantly impact its glycosylation pattern any more than the single D614G mutation does. This may be related to the positive selection pressure to maintain or enhance productive infectivity via engaging the host receptor, hACE2. We observed that N-glycans at N331 and N343 at the RBD remain highly processed while N234 and N165 also retain their Man_9_ and Man_5_ structures, respectively, consistent with glycosylation of these few sites playing active structural roles in modulating the conformational dynamics of the spike RBD^24, 25^. In contrast, both the unique ΔY144 and Δ69–70 deletions in the NTD of Alpha variant have been associated with immune escape from potent neutralizing antibodies^36–38^ against the single immunogenic supersite formed primarily by the disordered N1-N5 loops and bordered by glycans at N17, N74, N122, and N149^33, 39^. Glycans at N74 and N149 could not be visualized in our cryo-EM density map, consistent with their location on fairly flexible loop, unhindered from access and carry fully processed complex type structures. N122, on the other hand, is located nearby the N3 loop (residues 141-156) that is visibly affected by ΔY144 in the Alpha variant (Fig. 4A). This may have significantly affected the accessibility of N122, and hence the shift in the glycosylation status observed. The more exposed complex type N-glycans at N122, along with those at N74, N149 and N282 would collectively present an assorted range of host cell dependent glyco-epitopes that contribute to engaging the host glycan-binding auxiliary receptors^26, 27, 40^ in facilitated infectivity.

While a more definitive functional consequence cannot be readily delineated here, our findings attest to the sensitivity of glycosylation to point and/or deletion mutations in the primary sequence that may or may not lead to obvious conformational changes. More specifically, MS-based site-specific glycosylation mapping can detect these subtle alterations otherwise not informed by common protein structural analysis, at higher throughput and sensitivity once a standardized workflow is established.

## Acknowledgements

This work was supported by intramural funding of Academia Sinica (to STDH and KHK), an Academia Sinica Career Development Award (AS-CDA-109-L08 to STDH), and the Ministry of Science and Technology (MOST 109-3114-Y-001-001 to STDH). We thank the Academia Sinica Common Mass Spectrometry Facilities for Proteomics and Protein Modification Analysis (AS-CFII-108-107), and the Academia Sinica Cryo-EM Facility (AS-CFII-108-110) for data collection, both of which are funded by the Academia Sinica Core Facility and Innovative Instrument Project grants. We further thank the mammalian cell culture facility and the biophysics facility of Institute of Biological Chemistry, Academia Sinica, for supporting the protein production and characterizations, respectively. The technical support team of Protein Metrics is also gratefully acknowledged for helpful discussion in using the Byos software suite during the early stage of this work.

