## Supplemental Figure for "Distinct shifts in site-specific glycosylation pattern of SARS-CoV-2 spike proteins associated with arising mutations in the D614G and Alpha variants"

---

### Content

#### Supplementary Figures

Figure S1. Detailed heatmap of the quantifiable site-specific N-glycopeptides for each triplicate of the four different S-protein samples.

Figure S2. Zoomed-in views of the heatmap for quantifiable N-glycopeptides at select N-terminal domain sites, N74, N122 and N282.

Figure S3. The impact of applying different Byonic cut-off scores at and below 200 on the reliability of glycopeptide identification as evaluated by the number of matched peptide fragment ions.

Figure S4. Misassignment of glycopeptides attributable to missing glycan entries in the glycan library used in Byonic search.

Figure S5. Identification of terminal sulfated HexNAc and fucosylated $HexNAc_2$ on N-glycopeptides derived from recombinant trimeric S-proteins produced in HEK293F cells.

Figure S6. Purity and integrity of the intact trimeric SARS-CoV-2 spike proteins with and without furin cleavage site removed.

#### Supplementary Tables (available separately as Excel files)

**Table S1.** N-Glycan compositions included in the edited glycan library used for Byonic search.

**Table S2.** N-glycopeptides identified by Byonic search for each of the 12 S-protein sample datasets.

**Table S3.** Quantifiable site-specific N-glycopeptides for each of the four different S-protein samples.

**Table S4.** O-glycopeptides identified by Byonic search for each of the four different S-protein samples

---

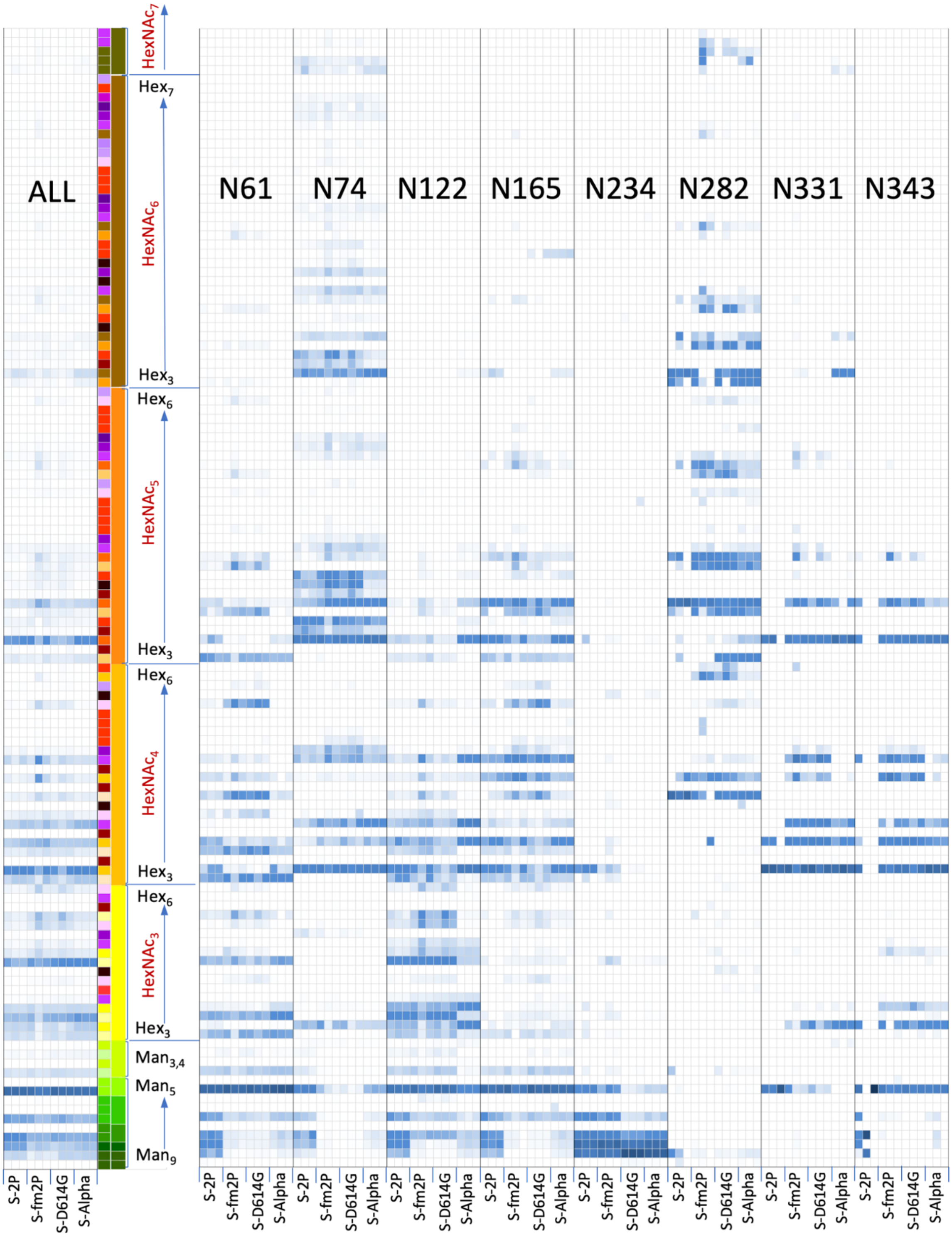


**Figure S1A. Detailed heatmap of the quantifiable site-specific N-glycopeptides for each triplicate of the four different S-protein samples.** Data were derived from Byos/Byologic output (Table S3) filtered according to the criteria defined and plotted directly without further correction or validation. The quantified peak areas for each of the 12 datasets are normalized to 100% and relative abundance is indicated in shades of blue (intensity correlates with abundance). Glycopeptide entries were sorted by their assigned glycan compositions, from high mannose $Man_{9\text{-}5}GlcNAc_2$, to pauci mannose $Man_{3,4}GlcNAc_2$±Fuc, then increasing number of $HexNAc_{3\text{-}7}$ coupled with increasing number of Hex, with 0-1 Fuc, from bottom to top, using the same color codes defined in Fig 2. A simplified version would ignore additional Fuc, NeuAc and sulfate while a more complex version would take that into account, with those carrying $NeuAc_{1\text{-}3}$, sulfate and/or $Fuc_{2,3}$ accorded a distinct color. In this Figure, the values for all sites, namely summing the quantifiable glycopeptides from all sites that carry the same glycan composition, are also shown, along with those on specific N-glycosylation sites, N61, N74, N122, N165, N234, N282, N331, N343, from NTD and RBD.

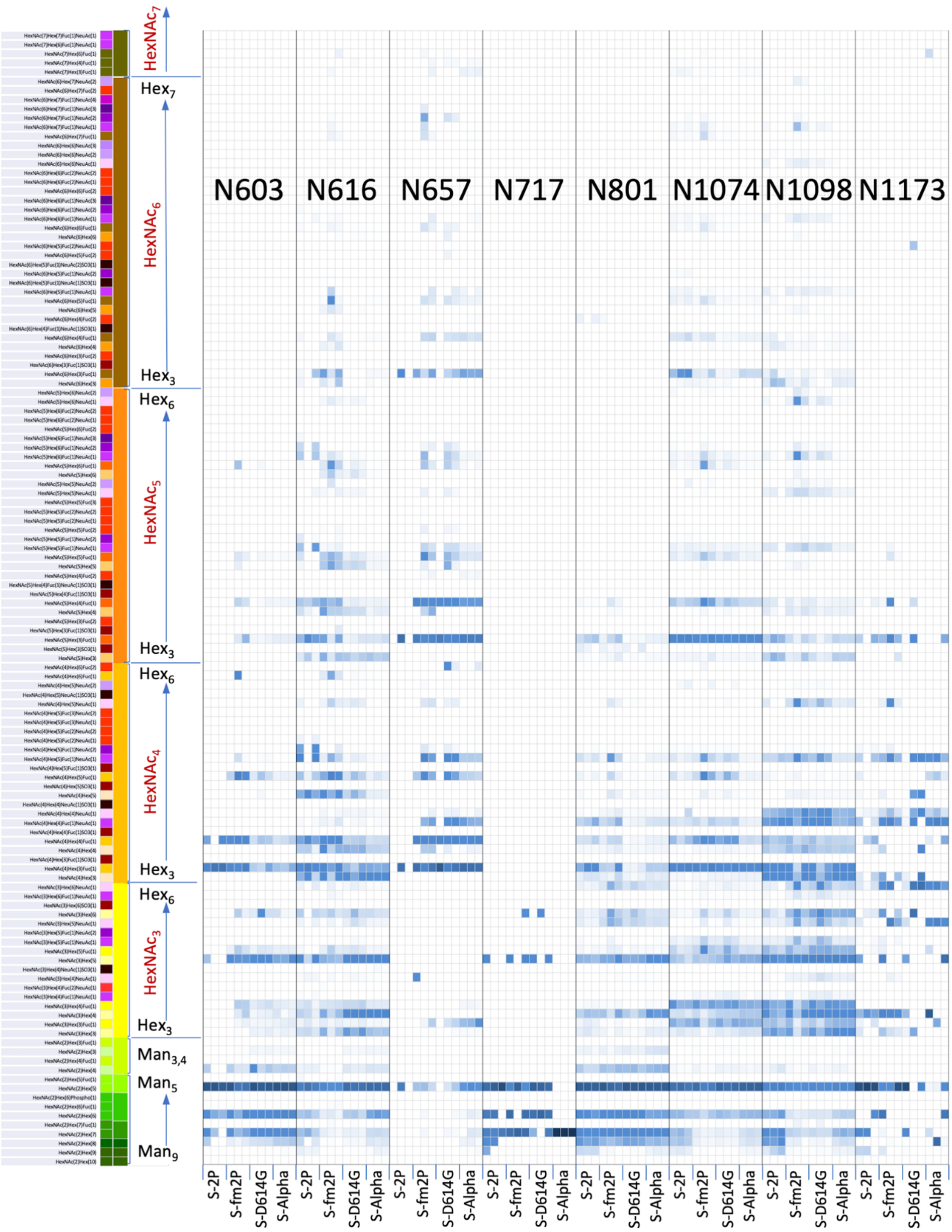


**Figure S1B. Detailed heatmap of the quantifiable site-specific N-glycopeptides for each triplicate of the four different S-protein samples** (continued from Fig S1A). Data were derived from Byos/Byologic output (Table S3) filtered according to the criteria defined and plotted directly without further correction or validation. The quantified peak areas for each of the 12 datasets are normalized to 100% and relative abundance is indicated in shades of blue (intensity correlates with abundance). Glycopeptide entries were sorted by their assigned glycan compositions, from high mannose $Man_{9-5}GlcNAc_2$, to pauci mannose $Man_{3,4}GlcNAc_2$±Fuc, then increasing number of $HexNAc_{3-7}$ coupled with increasing number of Hex, with 0-1 Fuc, from bottom to top, using the same color codes defined in Fig 2. Their exact compositions are also provided on the left of the corresponding color code strips. This Figure is a continuation of Figure S1A, which shows portion of the heatmap corresponding to summed all sites, and those on specific N-glycosylation sites of NTD and RBD. Here, the heatmap for the rest of the sites identified are shown.

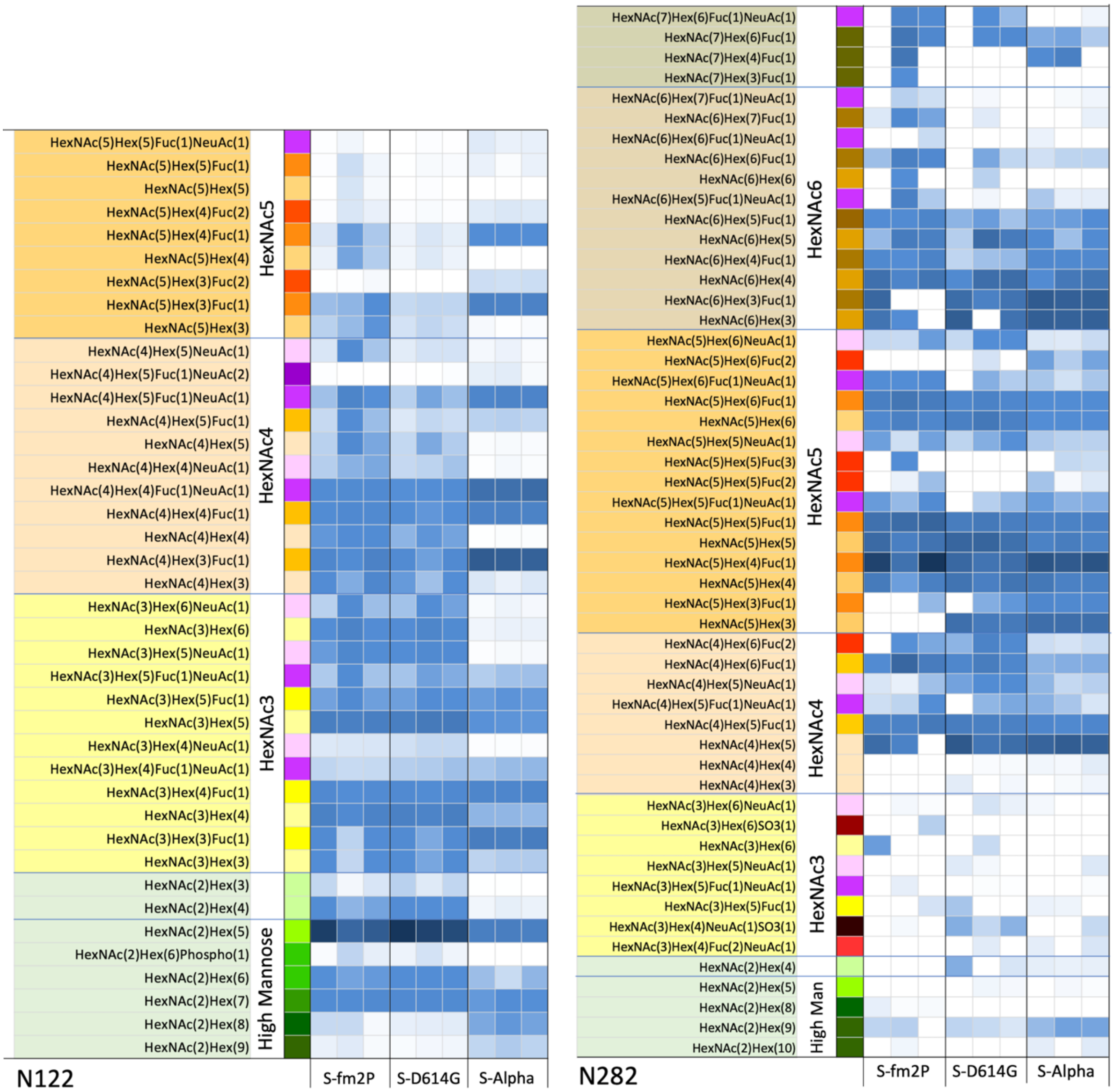


**Figure S2. Zoomed-in views of the heatmap for quantifiable N-glycopeptides at select N-terminal domain sites.** Shown here and continued onto next page are the individual heatmaps for three representative sites, N74, N122 and N282, that carry a full range of high mannose and complex type glycans. These are derived from the same dataset (Table S3) that are used to plot the full heatmap for all identified sites of all 4 S-protein samples in Fig S1. Only the data for each triplicate of the three S-protein samples, S-fm2P, S-D614G, and S-Alpha, are plotted. Data from S-2P are not as good quality due to overall lower abundance and hence less representative for a fair comparison. Glycopeptide entries with glycan compositions not found on each of the particular sites are removed in their corresponding heatmaps. Color codes are the same as that defined for Fig 2 and Fig S1. (Continue to next page for N74).

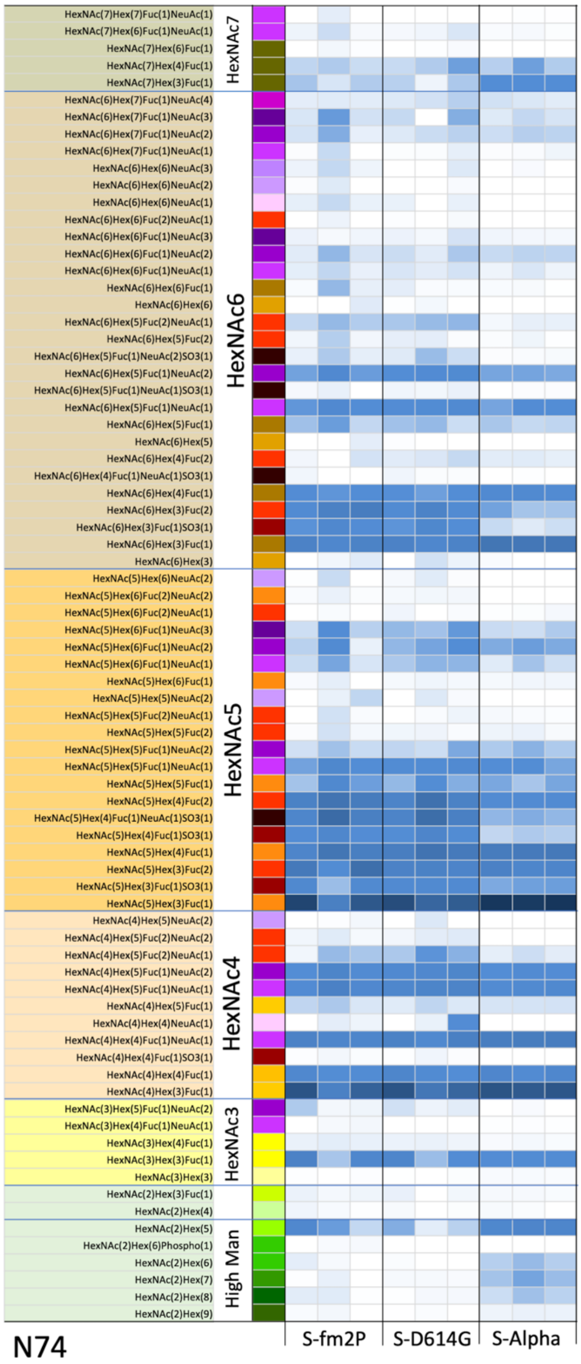


**Figure S2. Zoomed-in views of the heatmap for quantifiable N-glycopeptides at select N-terminal domain sites** (continue from previous page)**.** Shown here is the heatmap for site N74, for each triplicate of the three S-protein samples, S-fm2P, S-D614G, and S-Alpha.

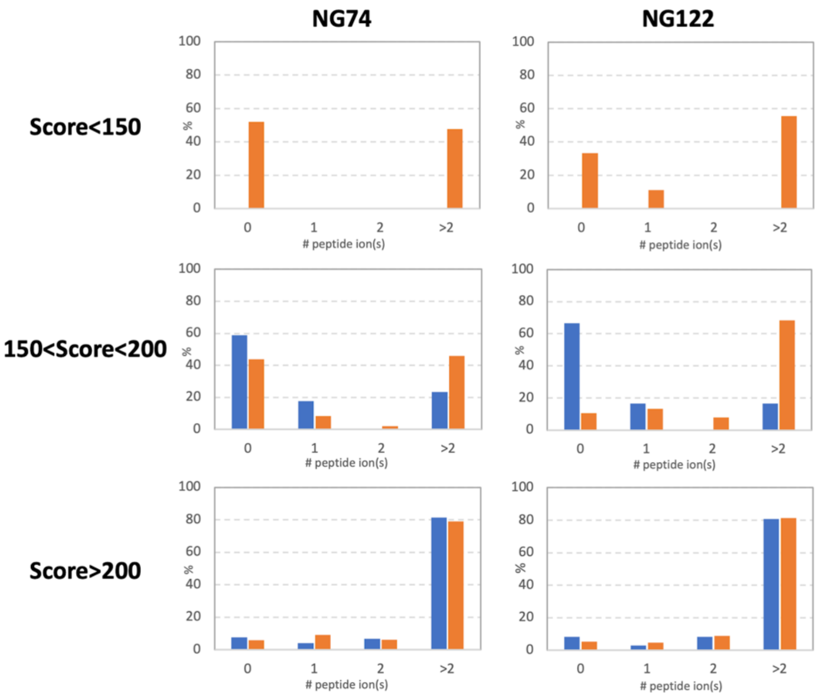


**Figure S3. The impact of applying different Byonic cut-off scores at and below 200 on the reliability of glycopeptide identification as evaluated by the number of matched peptide fragment ions**. HCD and EThcD $MS^2$ peptide spectrum matches (PSM) of glycopeptides at N74 (NG74, based on peptide FHAIHVSGTNGTK, HAIHVSGTNGTK, HAIHVSGTNGTKR) and N122 (NG122, based on LIVNNATNVVIK, IVNNATNVVIK) resulting from Byonic search were filtered first by PEP 2D< 0.001 and then different cut-off score at and below 200 were applied. PSMs satisfying both the PEP 2D and the indicated score cut-off were then grouped by the numbers of correctly assigned peptide b/c/y/z ion(s), using the datasets from S-2P (blue) and S-fm2P (orange) as examples.

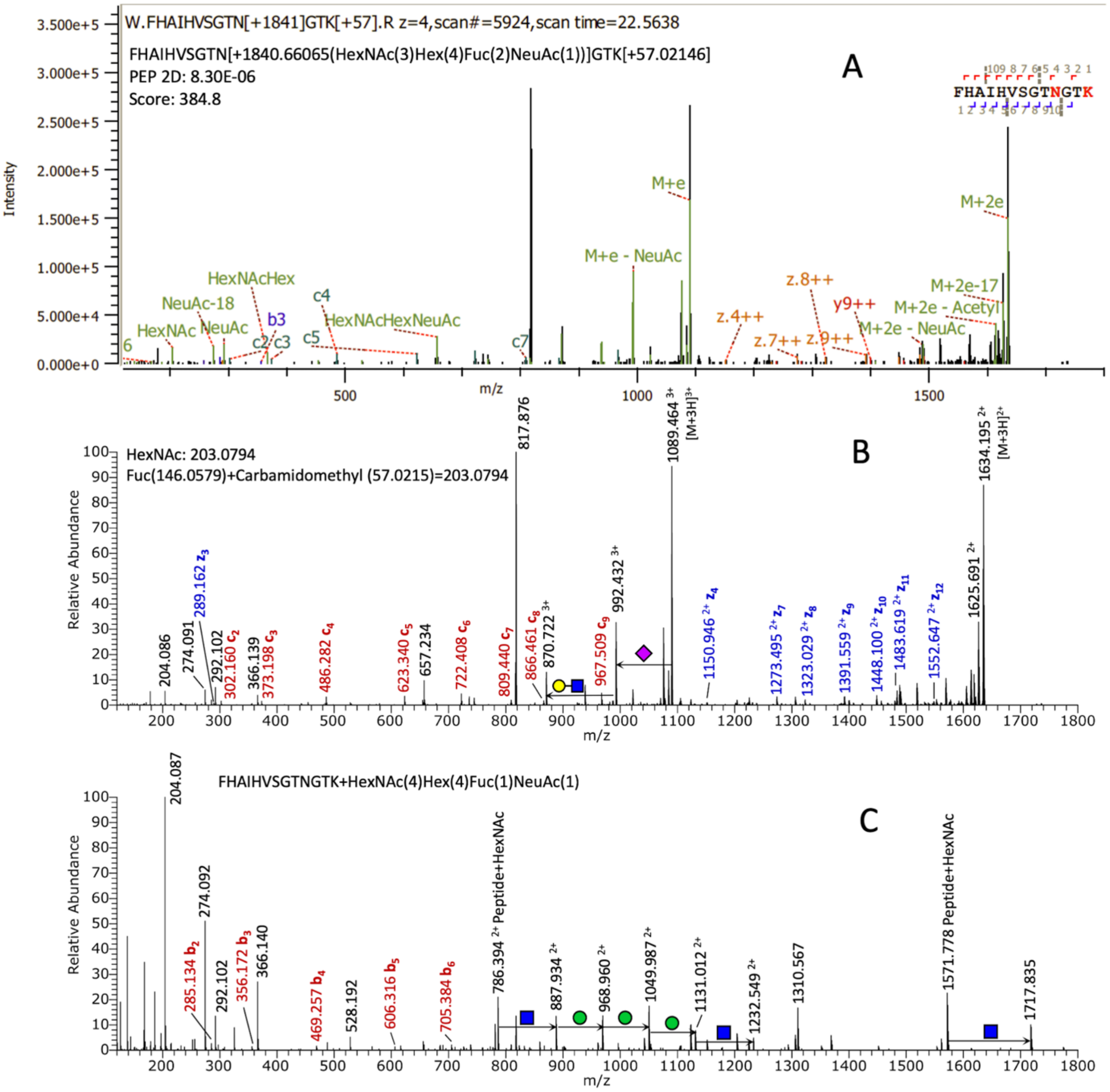


**Figure S4. Misassignment of glycopeptides attributable to missing glycan entries in the glycan library used in Byonic search**. (**A**) A glycopeptide at *m/z* 817.350 (z=4) was originally assigned as a peptide with carbamidomethyl modification on Lys and HexNAc(3)Hex(4)Fuc(2)NeuAc(1) on N74 by Byonic, based on its EThcD MS$^2$ spectrum. (**B**) Since the mass difference between one HexNAc (203.0794) and one Fuc (146.0579) is exactly the same as the mass increment of a carbamidomethyl modification (57.0215 u), Byonic has erroneously assigned a carbamidomethyl modification on the C-terminal Lys along with the N-glycan composition containing 3 HexNAc and 2 Fuc, instead of HexNAc(4)Hex(4)Fuc(1)NeuAc(1), which was not included the built-in 132 glycan library, to account for the extra 57 u. (**C**) Further manual inspection of the corresponding HCD MS$^2$ spectrum identified the presence of the Y1 (peptide backbone+HexNAc) ion at *m/z* 786.394 (z=2), corresponding to a peptide backbone without any carbamidomethyl modification, instead of *m/z* 814.905 if it carries one. Moreover, a carbamidomethylated Lys is not consistent with the $z_3$ ion detected at *m/z* 289.162 in the EThcD spectrum (**B**), which indicates a non-modified GTK. This ion was left unassigned by Byonic (**A**).

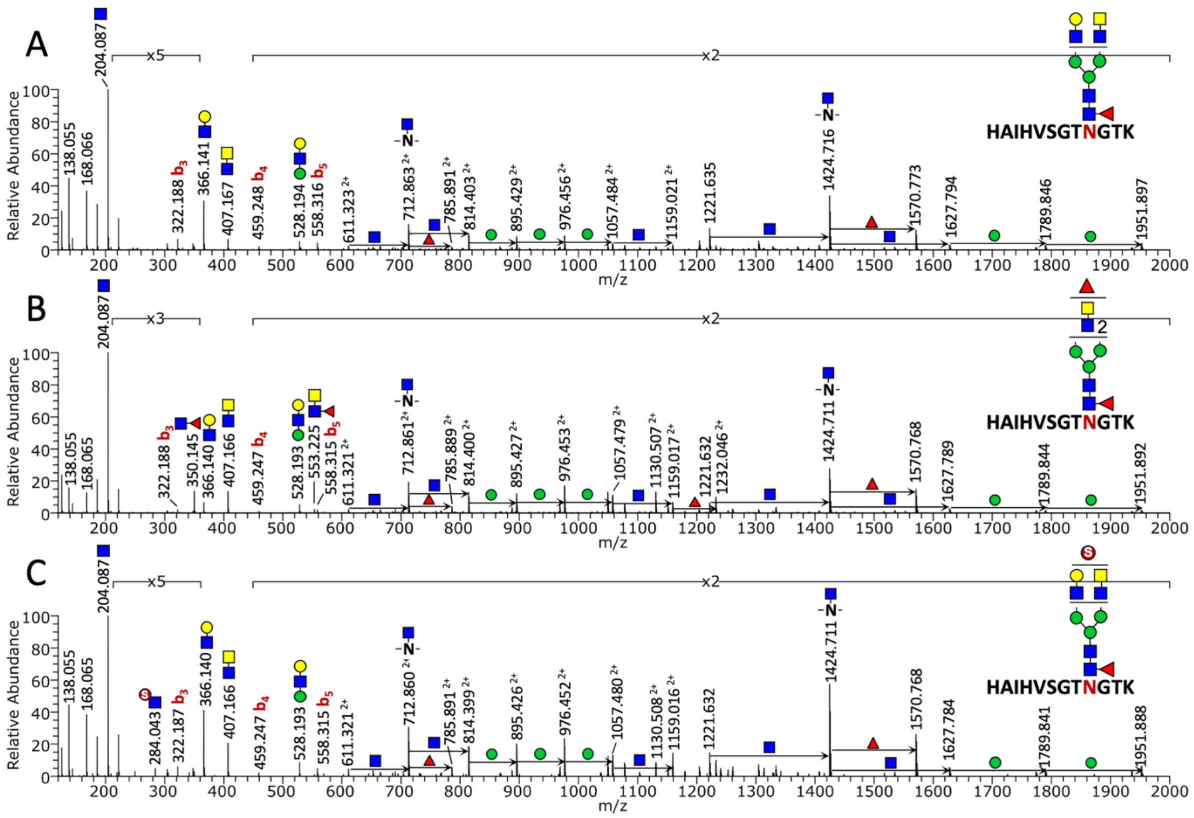


**Figure S5. Identification of terminal sulfated HexNAc and fucosylated $HexNAc_2$ on N-glycopeptides derived from recombinant trimeric S-proteins produced in HEK293F cells.** HCD $MS^2$ spectra of N74 glycopeptides sharing the same peptide backbone, HAIHVSGTNGTK, but carrying different glycans, detected at *m/z* 1011.103 (z=3) (**A**), 805.350 (z=4) (**B**), and 1037.756 (z=3) (**C**). The commonly detected oxonium ion at *m/z* 407.166 ($HexNAc_2^+$) is indicative of the presence of LacdiNAc modification. Additional $Fuc_1HexNAc_1^+$ and $Fuc_1HexNAc_2^+$ ions at *m/z* 350.145 and 553.225, respectively, identify the presence of a terminal $Fuc_1HexNAc_2$ glycotope, likely to be fucosylated LacdiNAc (**B**). Similarly, additional sulfated HexNAc ion at *m/z* 284.043 supports a sulfate modification on either the terminal LacNAc or LacdiNAc unit. The latter is more likely based on prior knowledge of the glycomic potentials of HEK293.

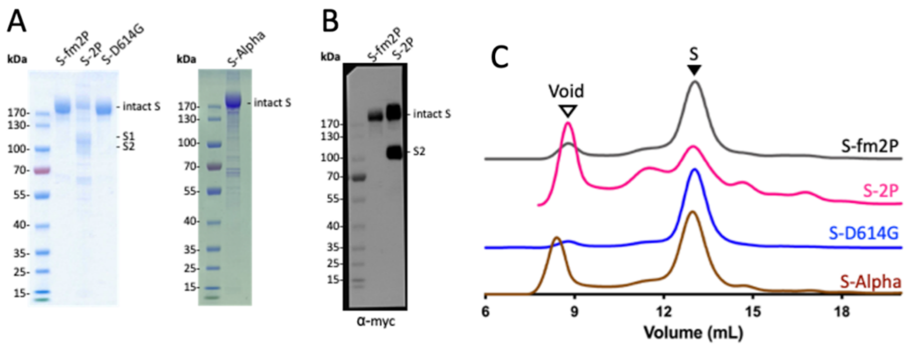


**Figure S6. Purity and integrity of the intact trimeric SARS-CoV-2 spike proteins with and without their furin cleavage site removed.** (**A**) SDS-PAGE analysis of the 4 different S proteins used in this study. Only S-2P, which retained its furin cleavage site yielded gel bands at lower molecular weight and identified by mass spectrometry analysis as cleaved S1 and S2 subunits. (**B**) Immunoblot analysis of the recovered S-fm2P and S-2P, detected by their C-terminal c-myc epitope tag and hence only the intact trimeric S-proteins and its S2 subunit would be visualized. (**C**) Size-exclusion chromatography profiles of S-fm2P (black), S-2P (magenta), S-D614G (blue) and S-Alpha (brown), with an elution volume (marked by solid arrow) consistent with their trimeric state.
